# DNA methylation correlates with transcriptional noise in response to elevated pCO_2_ in the eastern oyster (*Crassostrea virginica*)

**DOI:** 10.1101/2024.04.04.588108

**Authors:** Yaamini R. Venkataraman, Ariana S. Huffmyer, Samuel J. White, Alan Downey-Wall, Jill Ashey, Danielle M. Becker, Zachary Bengtsson, Hollie M. Putnam, Emma Strand, Javier A. Rodríguez-Casariego, Shelly A. Wanamaker, Kathleen E. Lotterhos, Steven B. Roberts

## Abstract

Ocean acidification significantly affects marine calcifiers like oysters, warranting the study of molecular mechanisms like DNA methylation that contribute to adaptive plasticity in response to environmental change. However, a consensus has not been reached on the extent to which methylation modules gene expression, and in turn plasticity, in marine invertebrates. In this study, we investigated the impact of pCO_2_ on gene expression and DNA methylation in the eastern oyster, *Crassostrea virginica*. After a 30-day exposure to control (572 ppm) or elevated pCO_2_ (2,827 ppm), whole genome bisulfite sequencing (WGBS) and RNA-Seq data were generated from adult female gonad tissue and male sperm samples. Although differentially methylated loci (DML) were identified in females (89) and males (2,916), there were no differentially expressed genes, and only one differentially expressed transcript in females. However, gene body methylation impacted other forms of gene activity in sperm, such as the maximum number of transcripts expressed per gene and changes in the predominant transcript expressed. Elevated pCO_2_ exposure increased gene expression variability (transcriptional noise) in males but decreased noise in females, suggesting a sex-specific role of methylation in gene expression regulation. Functional annotation of genes with changes in transcript-level expression or containing DML revealed several enriched biological processes potentially involved in elevated pCO_2_ response, including apoptotic pathways and signal transduction, as well as reproductive functions. Taken together, these results suggest that DNA methylation may regulate gene expression variability to maintain homeostasis in elevated pCO_2_ conditions and could play a key role in environmental resilience in marine invertebrates.

## Introduction

Ocean acidification (OA) is a pressing environmental issue resulting from the increased absorption of atmospheric carbon dioxide (CO_2_) by the world’s oceans [1]. Increased concentrations of oceanic CO_2_ leads to decreases in seawater pH, lower calcium carbonate concentrations, and reductions in aragonite saturation state [1]. These environmental alterations pose a significant threat to marine ecosystems, particularly to calcifying organisms such as oysters [2–5]. Reductions in saturation state can compromise calcification and increase the energetic cost of building shells [6–9], and can impact broad physiological pathways such as protein synthesis, energy production, metabolism, antioxidant responses, and reproduction [10–14]. Since oysters are vital for biodiversity, aquaculture, and coastal economies [15–17], understanding the biological responses of these organisms to OA is critical for assessing ecological impacts and developing strategies to mitigate negative effects of OA in oyster restoration and aquaculture production.

The ability of organisms to respond to and cope with OA are dependent on complex changes in whole-organism physiology, behavior, and molecular mechanisms [4,18–21]. Additionally, plasticity, or the capacity for an organism to adjust their physiology to maintain function and performance in response to environmental change [22], is another important factor governing responses to OA. Plastic responses to environmental stressors can also be inherited, providing a mechanism for the parent to pass on “environmental memory,” through mechanisms such as alterations in reproduction and offspring provisioning [23]. There are several examples of adult oyster exposure to OA yielding beneficial plasticity in offspring. For example, Sydney rock oysters (*Saccostrea glomerata*) exposed to high pCO_2_ during reproductive conditioning yielded larger and faster developing larvae compared to counterparts in ambient conditions [24,25], Olympia oyster (*Ostrea lurida*) offspring with parental history of OA exposure had better performance than ambient counterparts when outplanted [26], and exposure of adult eastern oysters (*Crassostrea virginica*) to elevated pCO_2_ increased larval growth rates [27]. However, parental exposure to OA in the Pacific oyster (*Crassostrea gigas*), northern quahog (*Mercenaria mercenaria*), and bay scallop (*Argopecten irradians*) led to poorer larval survival [28,29]. These contrasting results demonstrate that the capacity for intergenerational plasticity is complex, and requires additional research. As adult reproductive tissue gives rise to gametes and offspring, it is critical to examine the molecular mechanisms impacting reproductive homeostasis in response to OA.

Epigenetic modifications are a potential mediator of transgenerational plastic responses to environmental change. These modifications alter regulation of gene expression without altering the DNA sequence itself, and include multiple interacting mechanisms such as DNA methylation, chromatin remodeling, histone modifications, and non-coding RNAs [30]. DNA methylation — the addition of a methyl group to the cytosine adjacent to a guanine in a CpG dinucleotide [30,31] — is the most studied epigenetic mechanism in marine invertebrates [32]. DNA methylation can change expression of genes by altering binding of transcriptional machinery and can affect variability of transcription [31]. Changes in methylation are thought to be associated with phenotypic changes, with several examples of this connection illustrated in mammalian literature [33]. Several studies in marine invertebrates have also demonstrated the potential for methylomes (i.e. information of DNA methylation in a genome) to be inherited by offspring [34–38]. Methylomes are responsive to OA across a wide variety of taxa, such as pteropods [39], copepods [36,37], corals [40], sea urchins [41], and oysters [42–46]. These methylation responses are often species-specific, even within the same taxa [33]. For example, genes involved in protein ubiquitination, cytoskeletal regulation, and metabolism were all differentially methylated in various tissues and species, including the mantle [42] and gonad [44] of *C. virginica*, the mantle [45] and larvae [43,47] of the Hong kong oyster (*Crassostrea hongkongensis*), and the reproductive tissue [46] of female *C. gigas.* Genes with differential methylation were not necessarily orthologous across species. However, common processes impacted by DNA methylation across species and tissue types suggest a role for methylation in molluscan OA response.

Even though methylation is environmentally-responsive, direct connections between methylation and plasticity are not readily apparent in molluscs. Where gene expression has been used as a proxy for plasticity, molluscan studies report no relationship between differential gene expression and genes with differential methylation in response to environmental stress [34,42,45,48,49]. One exception in molluscan literature is Dang et al. [47], which found significant overlap between differentially expressed genes and differentially methylated genes in juvenile *C. hongkongensis* after exposure to OA conditions as larvae. Differentially expressed and methylated genes were often methylated in introns, and had roles in signal transduction [47]. Studies that do not explore gene expression and methylation have found associations between methylation and phenotype, although causal relationships have not been established. While differences in mantle methylomes cannot be linked to changes in calcification [42,45], there is evidence associating differential methylation with the ability to reproduce in stressful OA conditions [46], which in turn could be linked to maladaptive plasticity in larval *C. gigas* [29].

Additionally, differential methylation of *C. hongkongensis* larvae exposed to elevated pCO_2_ was associated with higher metamorphosis rates but poor substratum selection [43]. Taken together, this suggests that environmentally-responsive DNA methylation may play a role in modulating phenotypic plasticity in adult reproductive tissue, and could contribute to intergenerational carryover effects. Therefore, it is important to investigate knowledge gaps surrounding the role of methylation in altering gene expression and plasticity in oysters exposed to OA during reproductive conditioning.

This study explored the impact of OA on gene expression and DNA methylation in *C. virginica* adult reproductive tissue, and the relationship between gene expression and methylation (**Figure 1**). We used oysters from an accompanying study, which found that parental exposure to elevated pCO_2_ increased growth rates in larvae, especially when those larvae were also reared in elevated pCO_2_ conditions [27]. Female oysters in control and elevated pCO_2_ conditions did not have any differences in egg quality or size, suggesting that epigenetic mechanisms may have mediated the observed carryover effect [27]. We build on previous work demonstrating environmentally-sensitive methylation in *Crassostrea* spp. reproductive tissue [44,46] by examining sex-specific effects of OA on reproductive tissue responses. As female and male oysters undergo distinct gametogenic processes, and egg and sperm have unique roles in reproduction, it follows that there are sex-specific methylation [50–52] and gene expression [53] patterns in reproductive tissue that would confound findings if analyzed together. We compared female gonad tissue and male sperm exposed to control and elevated pCO_2_ conditions to investigate 1) the influence of genetic variation on methylation and gene expression; 2) the impact of OA on changes in gene activity and methylation; and 3) the functional role of DNA methylation in regulating gene activity. We define gene activity as encompassing gene-level expression, but also transcript-level processes such as transcript expression, changes in the number of unique transcripts expressed per gene, shifts in the predominant isoform expressed, and alternative splicing. By incorporating transcript-level analyses, we can further elucidate the functional role of methylation in regulating gene expression, and therefore plasticity, in response to environmental change. Through this comprehensive approach, we seek to uncover the molecular basis of oyster responses to OA, providing insights into potential impacts on reproduction and the molecular mechanisms involved in marine invertebrate stress response.

**Figure 1.**
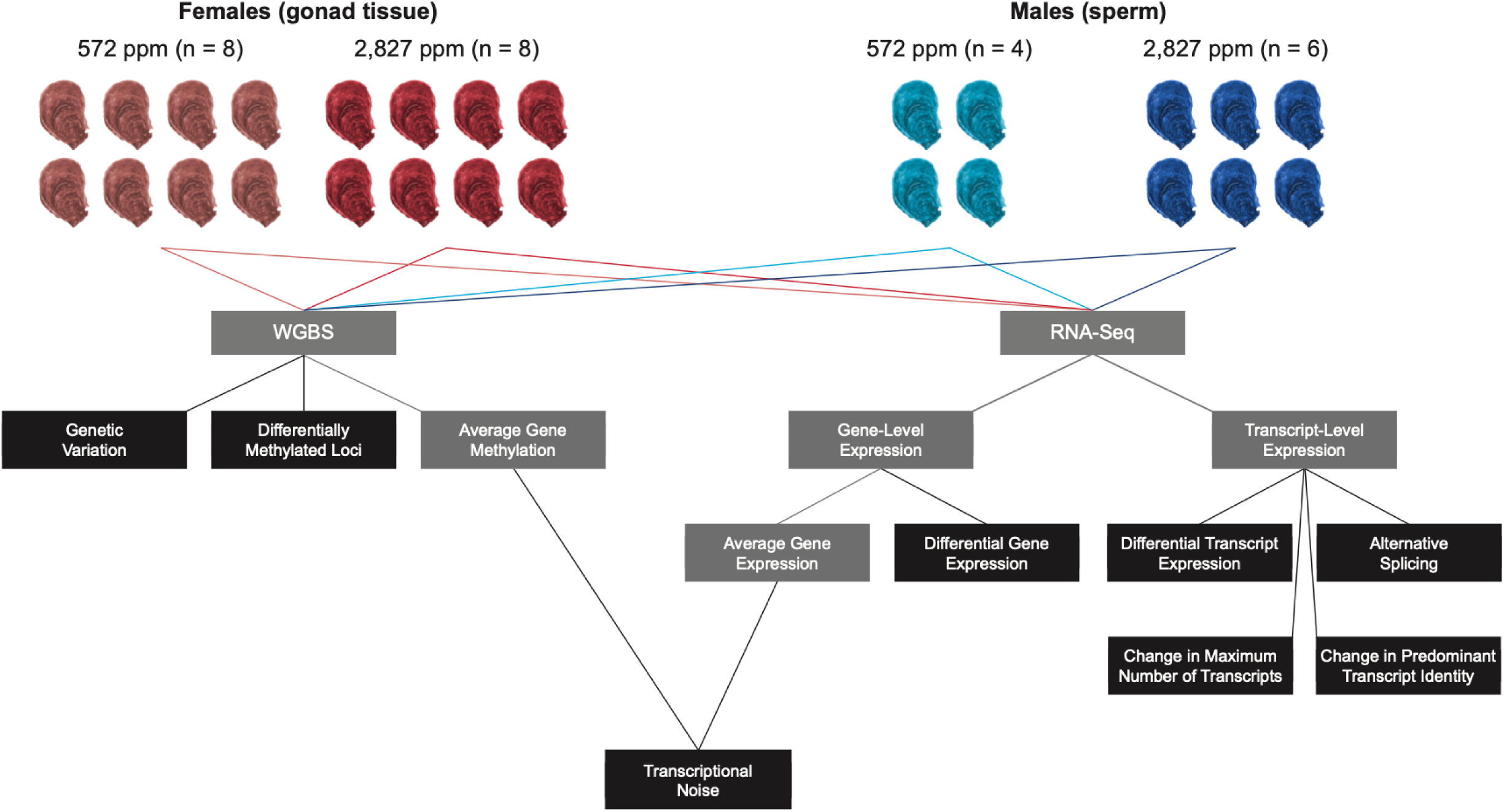
Conceptual diagram of study design. Female gonad tissue (n = 16) and male sperm (n = 10) from oysters exposed to either control (572 ppm) or elevated pCO_2_ (2,827 ppm) were used for Whole Genome Bisulfite Sequencing (WGBS) and RNA-Seq analysis. Black boxes indicate results discussed in this manuscript.

## Results

### Seawater Chemistry

As previously reported in McNally et al. [27], seawater of control and elevated pCO_2_ conditions were 572 ppm and 2827 ppm, respectively. The Ω_aragonite_ of seawater differed significantly between control and elevated pCO_2_ conditions (Welch’s 2-sample t-test: t = 12.33, df = 7, p-value < 0.0001).

### Genetic Variation

The full Whole Genome Bisulfite Sequencing (WGBS) dataset of all samples (female_control_ = 8, female_exposed_ = 8, male_control_ = 4, male_exposed_ = 6) was used to examine genetic variation, and the influence of that variation on gene methylation and expression. Even though the full dataset contains different cell types (*i.e.* female gonad tissue and male sperm), we wanted to determine the relatedness between female and male oysters. After filtering, we kept 2,343,637 single nucleotide polymorphisms (SNPs) out of 144,873,997 possible sites to generate a pairwise relatedness matrix and calculate genetic distance. Pairwise genetic, methylation, and gene expression distances were calculated using euclidean distances. There was no significant correlation between pairwise genetic distance and pairwise methylation distance (R = −0.16, *P*-value = 0.99) (**Supplementary File S1**) nor genetic distance and pairwise gene expression distance (R = −0.053, *P*-value = 0.76) (**Supplementary File S1**). Because pairwise genetic distance between samples was not related to pairwise distance in methylation nor gene expression between samples, genetic distance was not incorporated in downstream analyses of methylation and gene expression.

### Gene Activity

#### Gene and Transcript Expression

Across all female and male samples, 67.57% of RNA reads aligned to the *C. virginica* genome. Differentially expressed genes (DEG) and differentially expressed transcripts (DET) were identified between control and OA-exposed samples in female gonad tissue and male sperm samples separately. There were no DEG in females, but a single DET was identified (**Supplementary File S2**). The transcript was associated with gene ID LOC111134855 (uncharacterized). No DEG nor DET were found in males.

Changes in the maximum number of transcripts expressed per gene between control and OA-exposed samples were characterized as another measure of gene activity. In female gonads, a total of 3,966 genes (about 10% of total expressed genes) showed differences in the maximum number of transcripts expressed between female control and OA-exposed samples (**Table 1**). Under OA conditions in female gonads, 1,529 (38.6%) genes had a greater number of transcripts expressed per gene, while 2,437 had fewer transcripts expressed per gene (**Table 1**; **Supplementary File S3**). A total of 211 biological processes were enriched in genes with more transcripts under elevated pCO_2_, with the top ten significantly enriched GO terms including establishment of localization (*P*-value = 2.0 x 10^-10^) and cell adhesion (*P*-value = 2.6 x 10^-10^). Genes with less transcripts expressed under elevated pCO_2_ conditions had 469 enriched biological processes. The top ten significantly enriched GO terms included anatomical structure development (*P*-value = 5.3 x 10^-14^) and signaling (*P*-value = 7.0 x 10^-13^). Statistical output for all GO terms can be found in **Supplementary File S4**.

**Table 1.** Gene and transcript expression patterns attributable to elevated pCO_2_ conditions. Changes in maximum number of transcripts are described with respect to elevated pCO_2_ exposure, with increases in maximum number of transcripts representing more transcripts expressed in elevated pCO_2_-exposed samples, and decreases representing more transcripts expressed in control samples. Genes with multiple (≥ 2 transcripts) were considered for predominant transcript analyses.

| Sex | Total Genes Expressed | Genes with More Transcripts Expressed | Genes with Less Transcripts Expressed | Genes with $\geq 2$ Transcripts | Genes with a Shift in Predominant Transcript |
| --- | --- | --- | --- | --- | --- |
| Female | 39,047 | 1,529 | 2,437 | 38,575 | 2,252 |
| Male | 39,299 | 3,123 | 1,204 | 10,559 | 1,768 |

In sperm, 3,123 (72.2%) of genes that exhibited changes in the maximum number of transcripts had a greater number of transcripts expressed between control and OA-exposed samples, while 1,204 (27.8%) had fewer transcripts expressed in elevated pCO_2_ conditions (**Table 1**; **Supplementary File S3**). Genes with more transcripts expressed in elevated pCO_2_ conditions in sperm had 391 overrepresented biological processes, with the top ten significantly enriched GO terms including signaling (*P*-value = 4.9 x 10^-9^) and cytoskeletal organization (*P*-value = 1.9 x 10^-8^) (**Supplementary File S4**). Genes with fewer transcripts expressed in elevated pCO_2_ conditions had 238 enriched biological processes, with the top ten significantly enriched GO terms including organelle organization (*P*-value = 4.0 x 10^-7^) and intracellular signal transduction (*P*-value = 1.6 x 10^-6^) (**Supplementary File S4**).

### Predominant Transcript Identification

Low pH-induced shifts in the predominant transcript expression were assessed separately for female reproductive tissue and male sperm, where the predominant transcript was defined as the most highly expressed transcript within a gene. Of the 38,575 genes that had expression data for multiple transcripts in female gonads, 2,252 genes (5.83%) had shifts in the predominant transcript (**Table 1**; **Supplementary File S3**). A total of 226 biological process GO terms were overrepresented in genes where the predominant transcript shifted, with the top ten significantly enriched GO terms including regulation of small GTPase mediated signaling (*P*-value = 6.2 x 10^-9^), regulation of signal transduction (*P*-value = 5 x 10^-6^), and reproduction (*P*-value = 8.1 x 10^-6^). Statistical output for all GO terms can be found in **Supplementary File S4**.

For sperm, 1,768 (16.7%) out of 10,559 genes with data for multiple transcripts had shifts in the predominant transcript (**Table 1**; **Supplementary File S3**). Thirty-three biological processes were enriched in genes with shifts in the predominant transcript. The top 10 significantly enriched terms included actomyosin structure organization (*P*-value = 0.0015) and chemotaxis (*P*-value = 0.0017) (**Supplementary File S4**).

### Alternative Splicing

For each sex, we described patterns of alternative splicing by testing for variation in expression across exon position and treatment using ANOVA-simultaneous components analyses (ASCA) [54], calculated as the expression of exons 2 through 6 relative to expression of the first exon for every gene in each sample. ASCA analyses identified groups of genes with shared expression patterns across exons (e.g., genes that have highest expression of exon 1 relative to other exons, genes that have highest expression of exons 1 and 2 relative to other exons, etc.). Principal components (PC) of the ASCA model describe variation in patterns of gene expression across exons. Of the ten PC identified by the ASCA model for each sex, PC 1-6 explained greater than 1% of the variation in relative exon expression, and were selected for further analysis (**Supplementary File S5**). In both females (**Supplementary File S6**) and males (**Supplementary File S7**), PC 1-4 described alternative splicing patterns that did not differ by treatment, and explained the majority of variation in alternative splicing patterns (91.77% in females, 86.72% in males) (**Supplementary File S5**). In contrast, PC 5-6 described alternative splicing patterns that differed between treatments in females (**Figure 2)** and males (**Figure 3)**. Patterns of alternative splicing described by PC 5 and PC 6 accounted for low variance relative to PC 1-4 (7.69% in females, 10.89% in males; **Supplementary File S5**). Due to the low variance explained by PC 6 specifically (approximately 1.5%), confidence intervals could not be calculated for this pattern.

**Figure 2.**
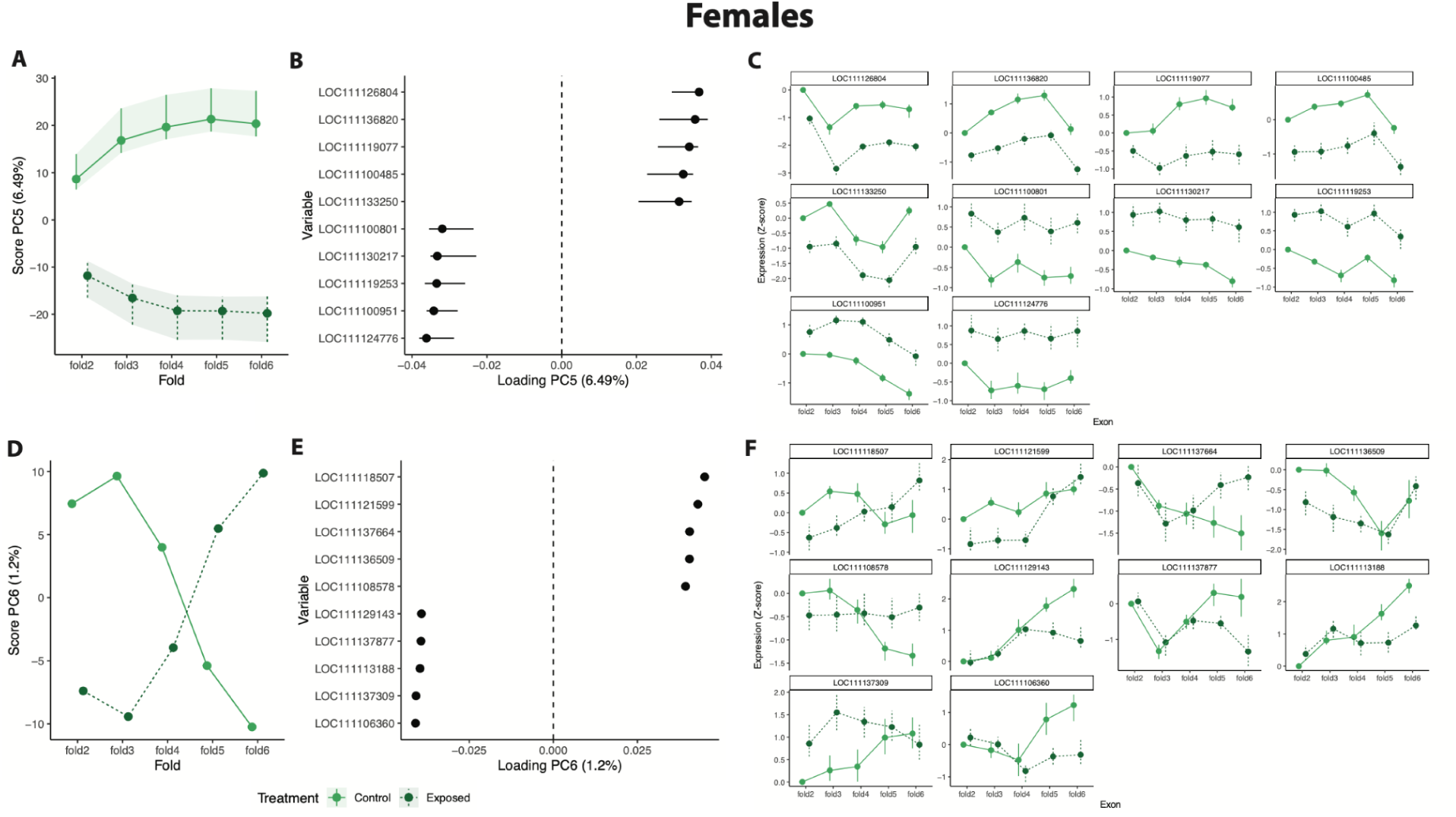
Principal components identified by ANOVA-simultaneous components analysis (ASCA) describing patterns of alternative splicing that show differential effects of treatment in females. A) PC 5 characterizes genes with differential expression in exons 2-6 relative to exon 1 between treatments. In all plots, dark green indicates exposed treatment and light green indicates control treatment. Shading indicates 95% confidence intervals. B) Top ten genes associated with alternative splicing patterns characterized by PC 5, ordered by PC loading score. Error bars indicate 95% confidence intervals with variables indicating genes. C) Expression (z-score) of exon position for each of the top ten genes associated with PC5. Panel title indicates gene. D) PC 6 characterizes genes with differential expression in the exons 2-4 relative to exon 1 or in exons 5-6 relative to exon 1 between treatments. Confidence intervals could not be calculated for PC 6 and are therefore not shown. E) Top ten genes associated with alternative splicing patterns characterized by PC 6, ordered by PC loading score. F) Expression (z-score) of exon position for each of the top ten genes associated with PC 6.

**Figure 3.**
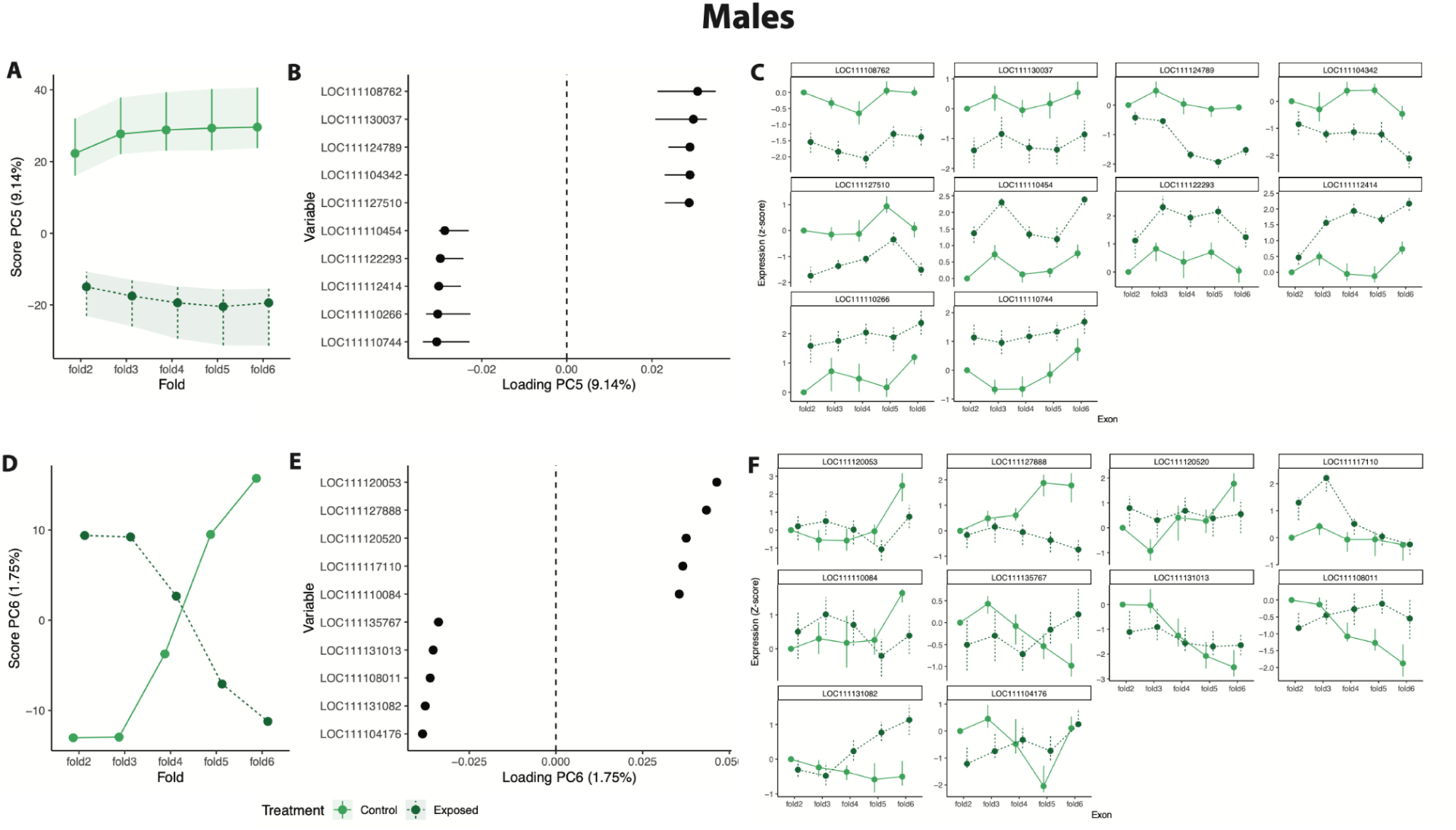
Principal components identified by ANOVA-simultaneous components analysis (ASCA) describing patterns of alternative splicing that show differential effects of treatment in males. A) PC 5 characterizes genes with differential expression in exons 2-6 relative to exon 1 between treatments. In all plots, dark green indicates exposed treatment and light green indicates control treatment. Shading indicates 95% confidence intervals. B) Top ten genes associated with alternative splicing patterns characterized by PC 5, ordered by PC loading score. Error bars indicate 95% confidence intervals with variables indicating genes. C) Expression (z-score) of exon position for each of the top ten genes associated with PC 5. Panel title indicates gene. D) PC 6 characterizes genes with differential expression in the exons 2-4 relative to exon 1 or in exons 5-6 relative to exon 1 between treatments. Confidence intervals could not be calculated for PC 6 and are therefore not shown. E) Top ten genes associated with alternative splicing patterns characterized by PC 6, ordered by PC loading score. F) Expression (z-score) of exon position for each of the top ten genes associated with PC 6.

There were four distinct patterns of alternative splicing that were not affected by treatment and were consistent between females **(Supplementary File S6)** and males **Supplementary File S7)**. Specifically, PC 1 described genes with expression that changes linearly across exon position, which includes genes that have both increasing and decreasing expression across exon position (females: **Supplementary File S6AB**, males: **Supplementary File S7AB**). PC 2 described genes that have either increased or decreased expression in middle exons (females: **Supplementary File S6CD**, males: **Supplementary File S7CD**) and PC 3 and PC 4 described genes that have either higher or lower expression in other combinations of exon positions (females: **Supplementary File S6E-H**, males: **Supplementary File S7E-H**). Since these patterns did not differ between treatments, we did not conduct functional enrichment analyses, and proceeded to analyze alternative splicing patterns that showed treatment differences (ie. PC 5 and PC 6).

There were two patterns of alternative splicing that revealed differences between treatments and were consistent in both females (**Figure 2**) and males (**Figure 3**). PC 5 described genes that show differential expression of exons 2-6 relative to exon 1 between treatments (**Figure 2A**, **Figure 3A**). We visualized the top ten genes that were most strongly associated with this pattern, revealing genes that show either lower or higher expression in exons 2-6 in the exposed treatment (**Figure 2BC**, **Figure 3BC**). PC 6 described genes that show differential expression of front and/or end exons relative to exon 1 (**Figure 2D**, **Figure 3D**). We also visualized the top ten genes associated with this pattern in females (**Figure 2F**) and males (**Figure 3F**), which revealed genes that had differential expression in exons 2-4 or exons 5-6 between treatments. Although the patterns were consistent across sexes, the top genes associated with these patterns differed.

We further analyzed the top 20 genes associated with PC 5 and PC 6. For females, five of the top 20 genes from PC 5 and PC 6 had changes in the maximum number of transcripts expressed (**Supplementary File S3**). There were 38 enriched biological process GO terms identified in the top female genes from PC5 and PC6. Statistical output for all GO terms can be found in **Supplementary File S4**. The top ten significantly enriched processes included protein UFMylation (ubiquitin-like modification; *P*-value = 6.1 x 10^-5^) and manganese ion transmembrane transport (*P*-value = 0.003) (**Supplementary File S4**). In males, four of the top 20 genes from PC 5 and PC 6 had changes in the maximum number of transcripts in males (**Supplementary File S3**). A total of 26 overrepresented biological processes were identified, with the top ten significantly enriched GO terms including regulation of metallopeptidase activity (*P*-value = 0.004) and acetyl-CoA metabolic process (*P*-value = 0.003).

### DNA Methylation Analysis

Approximately 57% of the quality trimmed reads were aligned to the genome, and on average 12.4% of the CpG loci were methylated. The average number of loci with a minimum of 10x coverage in the 26 individuals was 8.1M (56% of 14,458,703 total CpGs in the *C. virginica* genome). The baseline methylation landscape was characterized for each sex separately. A total of 12,678,572 and 11,991,497 non-C->T SNP CpGs had at least 10x coverage in at least one female or male sample, respectively. For both sexes, a majority of CpGs were lowly methylated (females: 10,212,421 CpGs, 80.5%; males: 9,657,277 CpGs, 80.5%), followed by highly methylated (females: 1,254,583 CpGs, 9.9%; males: 1,467,485 CpGs, 12.2%), and moderately methylated (females: 1,211,567 CpGs, 9.6%; males: 866,734 CpGs, 7.2%) (**Supplementary File S8**).

In females, highly methylated CpGs were found primarily in genes, with 662,082 CpGs (52.8%) in introns, 416,646 CpGs in CDS (33.2%), and 62,784 CpGs in exon UTR (5.0%) (**Supplementary File S8**). More highly methylated CpGs were located in the 1000 kb flanks downstream of genes (40,697 CpGs; 3.2%) than in flanks upstream of transcription start sites (7,896 CpGs; 0.6%). A total of 75,013 CpGs (6.0%) were found in intergenic regions, 80,887 CpGs (6.4%) in transposable elements, and 21,399 CpGs (1.7%) in lncRNA. Similarly, highly methylated CpGs in males were concentrated in genes, with more CpGs located in introns (812,892 CpGs, 55.4%) than CDS (438,908 CpGs, 30.0%) or exon UTR (77,872 CpGs, 5.3%) (**Supplementary File S8**). Downstream flanks contained more CpGs (46,650 CpGs, 3.2%) than upstream flanks (10,088 CpGs, 0.7%). Males also had higher concentrations of CpGs in transposable elements (93,876 CpGs, 6.4%), followed by intergenic regions (92,758 CpGs, 6.3%) and lncRNA (27,706 CpGs, 1.9%). The distribution of highly methylated CpGs in both sexes in genome features was significantly different than the distribution of all CpGs in the genome (chi-squared contingency tests; females: **Supplementary File S9**, males: **Supplementary File S10**).

### Differential Methylation at the CpG Locus-Level

There were 128 putative DML in female gonads and 4,175 putative DML in male sperm when comparing elevated pCO_2_-exposed and control oysters. After removing C->T SNPs, there were 89 and 2,916 female and male DML, respectively. Female DML were found predominantly in genes (74 DML; 83.1%), with 37 DML in introns (41.6%), 29 DML in CDS (32.6%), and 8 DML in exon UTR (9.0%) (**Figure 4**). Seven DML (7.9%) were found in intergenic regions, four DML (4.5%) were found in both upstream and downstream flanks, and one DML (1.1%) was found in both lncRNA and TE. The proportion of female DML found in either CDS or intergenic regions was significantly different from the proportion of 10x CpGs in those same genome features (chi-squared contingency test results can be found in **Supplementary File S11**). Gene ontology enrichment revealed 85 biological process GO terms enriched in genes containing DML The top ten significantly enriched processes included dolichol metabolic process and (*P*-value = 1.1 x 10^-5^) negative regulation of apoptotic process (*P*-value = 0.001). Statistical output for all GO terms can be found in **Supplementary File S4**.

**Figure 4.**
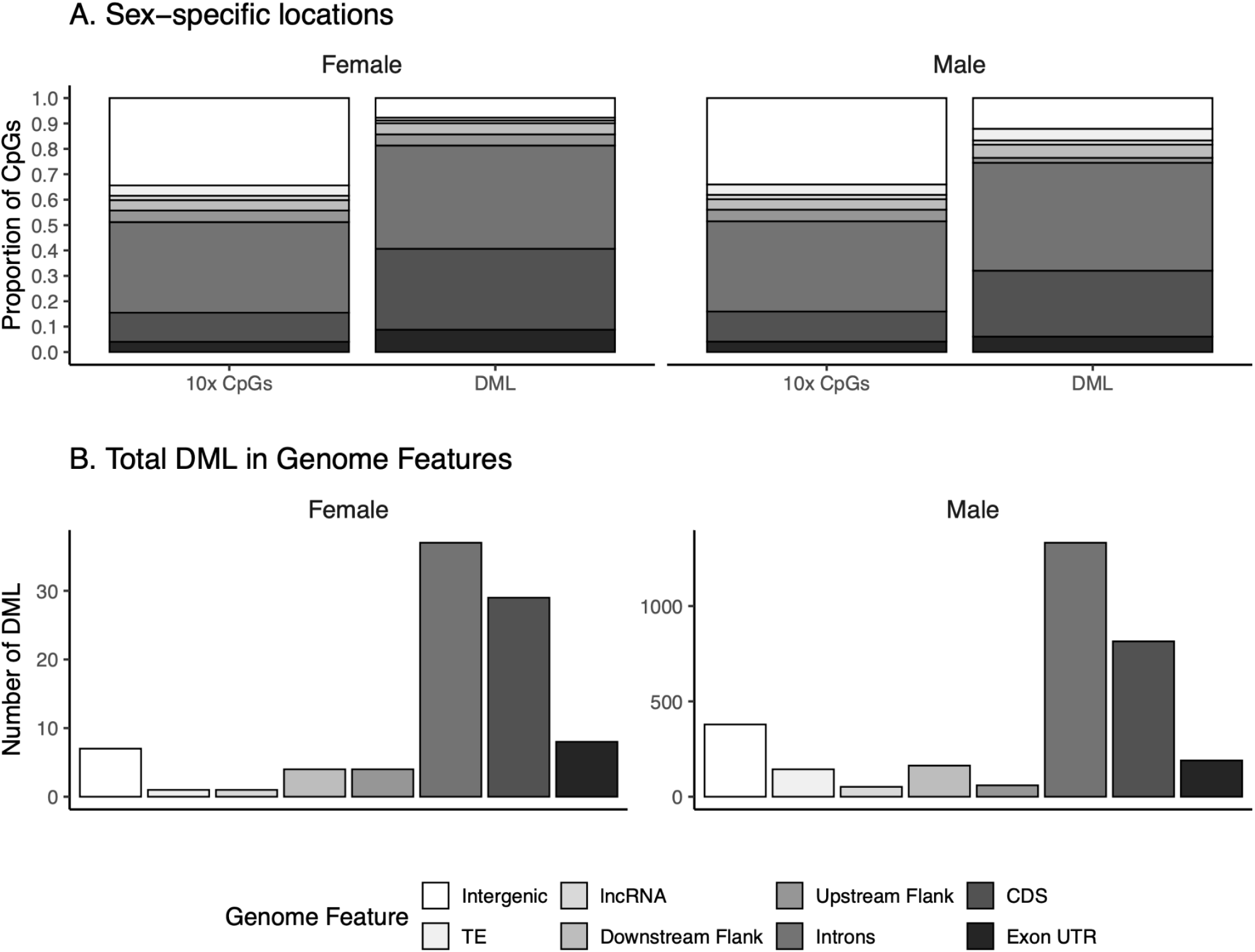
Genomic locations of 89 DML in female gonad tissue and 2,916 DML in male sperm. A) Proportion 10x CpGs with data versus DML in various genome features. B) Total number of DML in various genome features. All statistical test output comparing locations of DML with 10 CpGs can be found in **Supplementary File S11** for female gonads and in **Supplementary Files S12** for male sperm.

Similar to females, the majority of male DML were found in gene bodies (2,322 DML; 79.6%) (**Figure 4**). Within genes, 1,332 DML were found in introns (45.7%), 815 DML in CDS (27.9%), and 190 DML (6.5%) in exon UTR. More DML were found in downstream flanks (163 DML; 5.6%) than upstream flanks (60 DML; 2.1%). A total of 279 DML (9.6%) were found in intergenic regions, while 144 DML (4.9%) were in TE and 52 DML (1.8%) were in lncRNA. The proportion of male DML in exon UTR, CDS, introns, upstream flanks, downstream flanks, and intergenic regions differed significantly from the proportion of 10x CpGs in those genome features (chi-squared contingency test results can be found in **Supplementary File S12**). A total of 359 biological process GO terms were enriched in genes containing DML, with the top ten overrepresented processes including organelle organization (*P*-value = 7.1 x 10^-15^), regulation of GTPase activity (*P*-value = 1.2 x 10^-11^), and macromolecule localization (*P*-value = 9.8 x 10^-9^) (**Supplementary File S4**).

### Methylation Influence on Gene Activity

A significant and positive correlation was found between gene methylation and expression distance matrices, indicating that samples that were highly similar in gene expression were also highly similar in methylation (R = 0.843, *P*-value = 0.01) (**Supplementary File S1**). To investigate the reason for this higher-level correlation, methylation’s influence on gene activity was evaluated by examining its effect on 1) gene and transcript expression, 2) predominant transcript shifts, 3) transcriptional noise, and 4) alternative splicing.

### Influence of Methylation on Gene and Transcript Expression

The one transcript differentially expressed between control and elevated pCO_2_-exposed female oysters did not contain DML. Since a one-to-one relationship was not identified between DML and DET, the influence of methylation on other aspects of gene activity was explored. We first examined connections between average gene methylation and changes in the maximum number of transcripts expressed. The number of DML in genes with changes in the maximum number of expressed transcripts ranged from zero to two in females (**Supplementary File S13**) and zero to 25 in males (**Supplementary File S13**). A binomial model was used to evaluate the influence of the explanatory variable, gene body methylation, on whether or not there was a change in the response variable, the maximum number of transcripts expressed due to elevated pCO_2_ exposure. The number of DML in a gene, gene length, and gene expression were used as additional explanatory variables. The number of DML in a gene was not a significant predictor of a change in maximum transcripts expressed (**Table 2**, **Supplementary File S14**) in female gonad tissue or male sperm. Change in gene body methylation had a marginally significant negative effect on changes in the maximum number of transcripts male oysters (**Table 2**). For both sexes, change in gene expression and gene length were significant predictors (**Table 2**).

**Table 2.** Summary of binomial model for female and male oysters examining the influence of the explanatory variables of gene body methylation, DML count, gene length, and gene expression on the response variable of changes to the maximum number of transcripts expressed due to elevated pCO_2_ exposure. \**P*-value < 0.01; \*\**P*-value < 0.05.

| Explanatory Variable | Females |  |  | Males |  |  |
| --- | --- | --- | --- | --- | --- | --- |
| | $\beta$ Estimate | z-value | *P*-value | $\beta$ Estimate | z-value | *P*-value |
| Intercept | 0.24 | 0.67 | 0.51 | 2.66 | 6.88 | 6.00 x 10<sup>-12**</sup> |
| Change in Gene Body Methylation | 0.007 | 0.44 | 0.66 | -0.01 | -1.76 | 0.08* |
| DML Count | 0.35 | 0.23 | 0.82 | -0.05 | -1.07 | 0.29 |
| log<sub>2</sub>(Change in Gene Expression) | 0.65 | 11.31 | 1.13 x 10<sup>-29**</sup> | -0.54 | -5.86 | 4.63 x 10<sup>-9**</sup> |
| log<sub>10</sub>(Gene Length) | -0.19 | -2.14 | 0.03** | 0.18 | 6.27 | 3.67 x 10<sup>-10**</sup> |

### Influence of Methylation on Predominant Transcript Shifts

Several DML were found in genes with a shift in the predominant transcript expressed. In females, zero to three DML were found in these genes (**Supplementary File S13**), with zero to 25 DML in males (**Supplementary File S13**). Binomial models were used to quantify the impact of gene body methylation, DML count, gene expression, and gene length on whether or not there was a shift in the predominant transcript expressed due to low pH exposure. In female oysters, change in gene body methylation and number of DML in genes did not have a significant effect (**Table 3, Supplementary File S15**). However, change in gene body methylation had a significant positive effect on a shift in the predominant transcript in males (**Table 3, Supplementary File S15**). Gene expression and gene length were significant predictors of predominant transcript shift in both females and males (**Table 3**).

**Table 3.** Summary of binomial model for female and male oysters examining the influence of the explanatory variables of gene body methylation, DML count, gene length, and gene expression on the response variable, a shift in the predominant transcript due to elevated pCO_2_ exposure. \**P*-value < 0.01; \*\**P*-value < 0.05.

| Explanatory Variable | Females |  |  | Males |  |  |
| --- | --- | --- | --- | --- | --- | --- |
| | $\beta$ Estimate | z-value | *P*-value | $\beta$ Estimate | z-value | *P*-value |
| Intercept | -8.49 | -35.93 | $9.54 \times 10^{-283**}$ | -3.04 | -9.44 | $3.71 \times 10^{-21**}$ |
| Change in Gene Body Methylation | -0.005 | -0.58 | 0.56 | 0.01 | 2.03 | 0.04** |
| DML Count | -11.64 | -0.10 | 0.92 | 0.07 | 1.61 | 0.11 |
| log<sub>2</sub>(Change in Gene Expression) | -0.21 | -11.10 | $7.20 \times 10^{-140**}$ | 0.05 | 2.16 | 0.03** |
| log<sub>10</sub>(Gene Length) | 1.45 | 25.18 | $1.32 \times 10^{-28**}$ | 0.33 | 4.28 | $1.88 \times 10^{-5**}$ |

### Transcriptional Noise Analysis

Models demonstrate that both gene body methylation and expression have a significant influence on gene expression variability, or transcriptional noise (**Table 4**, **Figure 5**). All predictors were included in the best-fit model for each sex. For female gonads (model adjusted R^2^ = 0.60), transcriptional noise was negatively correlated with treatment (*P*-value = 0.002; **Figure 5A**), gene expression (*P-value* = 0; **Figure 5C**), gene body methylation (*P*-value = 0; **Figure 5E**), and the interaction between treatment and expression, but positively correlated with methylation variation (*P*-value = 1.87 x 10^-272^; **Figure 5G**), gene expression squared, the interaction between treatment and gene expression squared, and gene length (**Table 4**).

**Figure 5.**
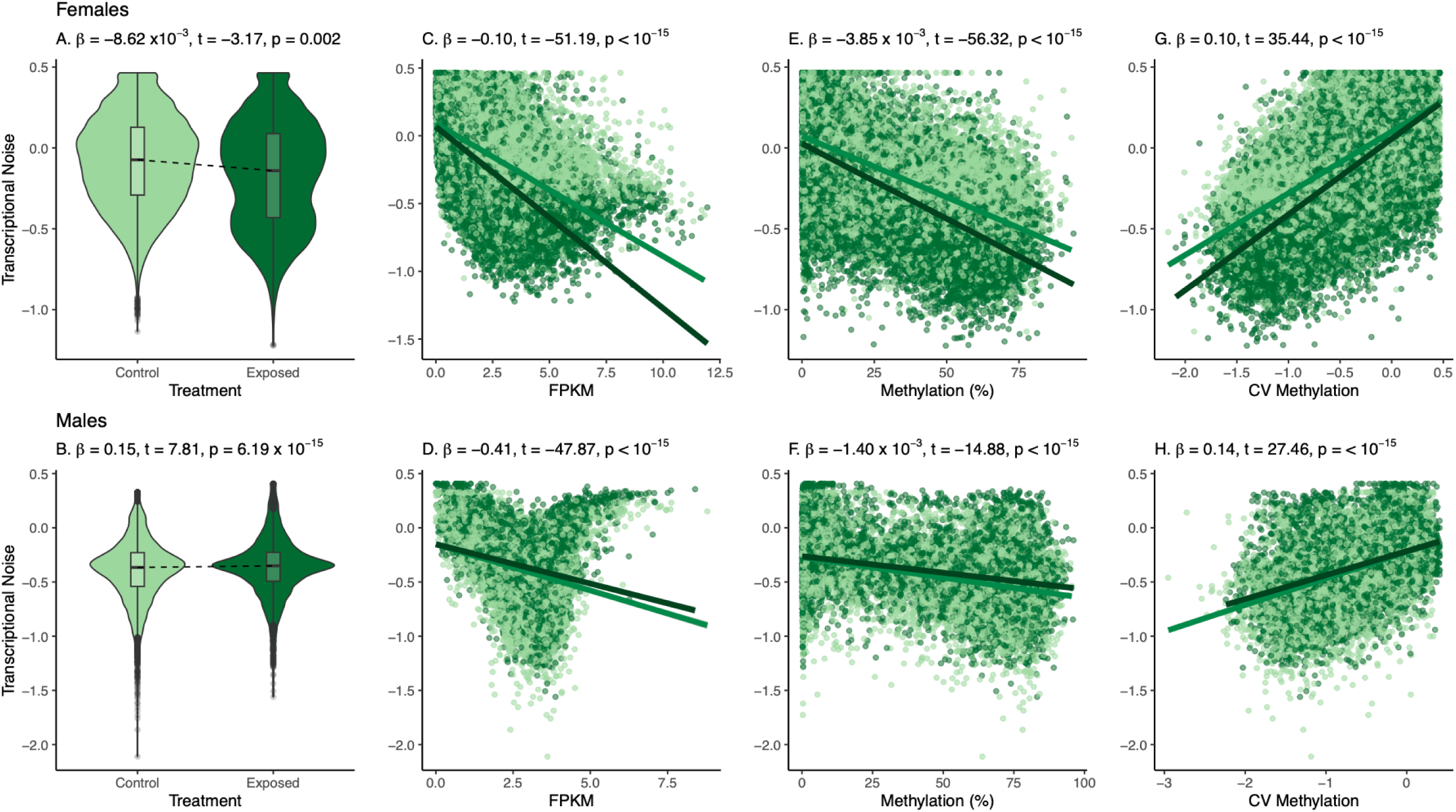
Relationship between transcriptional noise and A) and B) elevated pCO_2_ treatment; C) and D) FPKM; E) and F) gene methylation (%); and G) and H) CV of methylation as defined by the best-fit linear model for female gonads (adjusted R^2^ = 0.60) and male sperm (adjusted R^2^ = 0.38) oysters. Transcriptional noise and FPMK are log_10_ and log_2_ transformed, respectively, for visualization. Beta statistics, t-values, and *P*-values are included for each explanatory variable of the linear model. For panels C-H, each point represents values for a single gene, with associated control values in light green and elevated pCO_2_ values in dark green. Linear model fits are included for control and elevated pCO_2_ treatments. Additional model predictors, including polynomial terms for gene expression, can be found in **Table 3**.

**Table 4.**
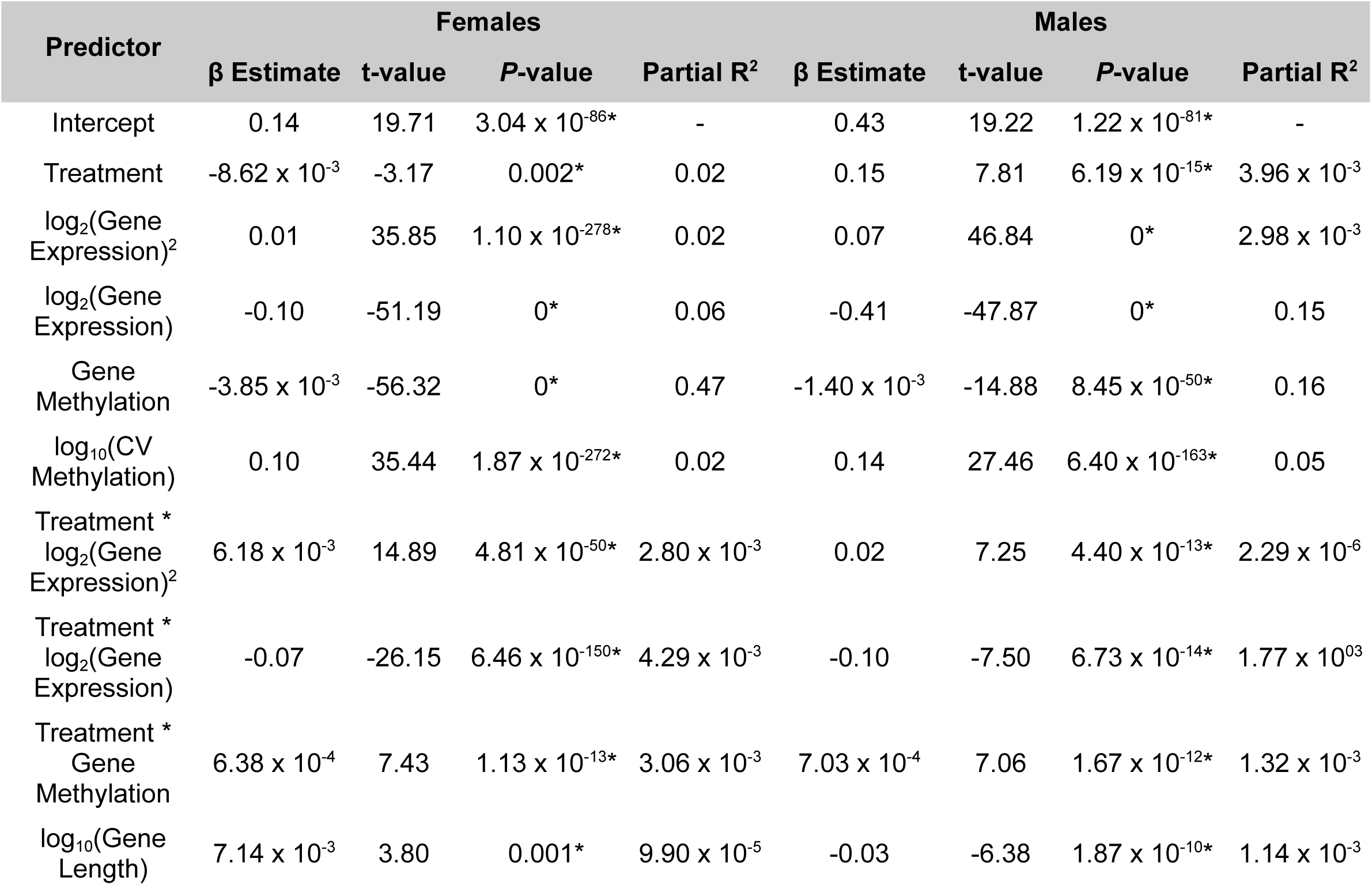
Summary of multiple linear regression examining the correlation between the explanatory variables of gene expression, gene body methylation, the CV of gene body methylation, gene length, and pCO_2_ treatment on the response variable of transcriptional noise, or the CV of gene expression. Separate models were constructed for female gonads (adjusted R^2^ = 0.60) and male sperm (adjusted R^2^ = 0.38). Coefficients of determination, or partial R^2^, were calculated separately for each predictor. \**P*-value < 0.05.

| Predictor | Females |  |  |  | Males |  |  |  |
| --- | --- | --- | --- | --- | --- | --- | --- | --- |
|  | β Estimate | t-value | *P*-value | Partial R<sup>2</sup> | β Estimate | t-value | *P*-value | Partial R<sup>2</sup> |
| Intercept | 0.14 | 19.71 | 3.04 x 10<sup>-86*</sup> | - | 0.43 | 19.22 | 1.22 x 10<sup>-81*</sup> | - |
| Treatment | -8.62 x 10<sup>-3</sup> | -3.17 | 0.002* | 0.02 | 0.15 | 7.81 | 6.19 x 10<sup>-15*</sup> | 3.96 x 10<sup>-3</sup> |
| log<sub>2</sub>(Gene Expression)<sup>2</sup> | 0.01 | 35.85 | 1.10 x 10<sup>-278*</sup> | 0.02 | 0.07 | 46.84 | 0* | 2.98 x 10<sup>-3</sup> |
| log<sub>2</sub>(Gene Expression) | -0.10 | -51.19 | 0* | 0.06 | -0.41 | -47.87 | 0* | 0.15 |
| Gene Methylation | -3.85 x 10<sup>-3</sup> | -56.32 | 0* | 0.47 | -1.40 x 10<sup>-3</sup> | -14.88 | 8.45 x 10<sup>-50*</sup> | 0.16 |
| log<sub>10</sub>(CV Methylation) | 0.10 | 35.44 | 1.87 x 10<sup>-272*</sup> | 0.02 | 0.14 | 27.46 | 6.40 x 10<sup>-163*</sup> | 0.05 |
| Treatment * log<sub>2</sub>(Gene Expression)<sup>2</sup> | 6.18 x 10<sup>-3</sup> | 14.89 | 4.81 x 10<sup>-50*</sup> | 2.80 x 10<sup>-3</sup> | 0.02 | 7.25 | 4.40 x 10<sup>-13*</sup> | 2.29 x 10<sup>-6</sup> |
| Treatment * log<sub>2</sub>(Gene Expression) | -0.07 | -26.15 | 6.46 x 10<sup>-150*</sup> | 4.29 x 10<sup>-3</sup> | -0.10 | -7.50 | 6.73 x 10<sup>-14*</sup> | 1.77 x 10<sup>03</sup> |
| Treatment * Gene Methylation | 6.38 x 10<sup>-4</sup> | 7.43 | 1.13 x 10<sup>-13*</sup> | 3.06 x 10<sup>-3</sup> | 7.03 x 10<sup>-4</sup> | 7.06 | 1.67 x 10<sup>-12*</sup> | 1.32 x 10<sup>-3</sup> |
| log<sub>10</sub>(Gene Length) | 7.14 x 10<sup>-3</sup> | 3.80 | 0.001* | 9.90 x 10<sup>-5</sup> | -0.03 | -6.38 | 1.87 x 10<sup>-10*</sup> | 1.14 x 10<sup>-3</sup> |

Inclusion of polynomial terms for gene expression reflects that gene expression variability is highest at very low and very high levels of gene expression. Methylation had the highest partial R^2^ (partial R^2^ = 0.47; **Table 4**), followed closely by model residuals (partial R^2^ = 0.40). For male sperm, transcriptional noise was positively correlated with treatment (*P*-value = 6.19 x 10^-15^; **Figure 5B**), negatively correlated with gene expression (*P*-value = 0; **Figure 5D**) and gene body methylation (*P*-value = 8.45 x 10^-50^; **Figure 5F**), and positively correlated with methylation variation (*P*-value = 6.40 x 10^-163^; **Figure 5H**). Transcriptional noise was also negatively correlated with the interaction between treatment and gene expression and gene length, but positively correlated with gene expression squared, the interaction between treatment and gene expression squared, and the interaction between treatment and gene body methylation (**Table 4**). The model residuals had the highest partial R^2^ (partial R^2^ = 0.62) followed by methylation (partial R^2^ = 0.16; **Table 4**). In both female gonads and male sperm, the statistically significant influence of pCO_2_ treatment on transcriptional noise was associated with a small effect size, likely due to inter-sample variability.

### Influence of Methylation on Alternative Splicing

The influence of methylation was examined in the top 20 genes from PC 5 and PC 6, which exhibited changes in alternative splicing from elevated pCO_2_ exposure. There were no DML found in top female gonad genes, while six male sperm genes contained DML. Average gene body methylation in control and exposed samples were investigated as predictors of alternative splicing using sex-specific binomial GLM. In both sexes, average gene body methylation in control samples, methylation in exposed samples, and gene length had no significant impact on alternative splicing status (**Table 5**; **Figure 6**).

**Figure 6.**
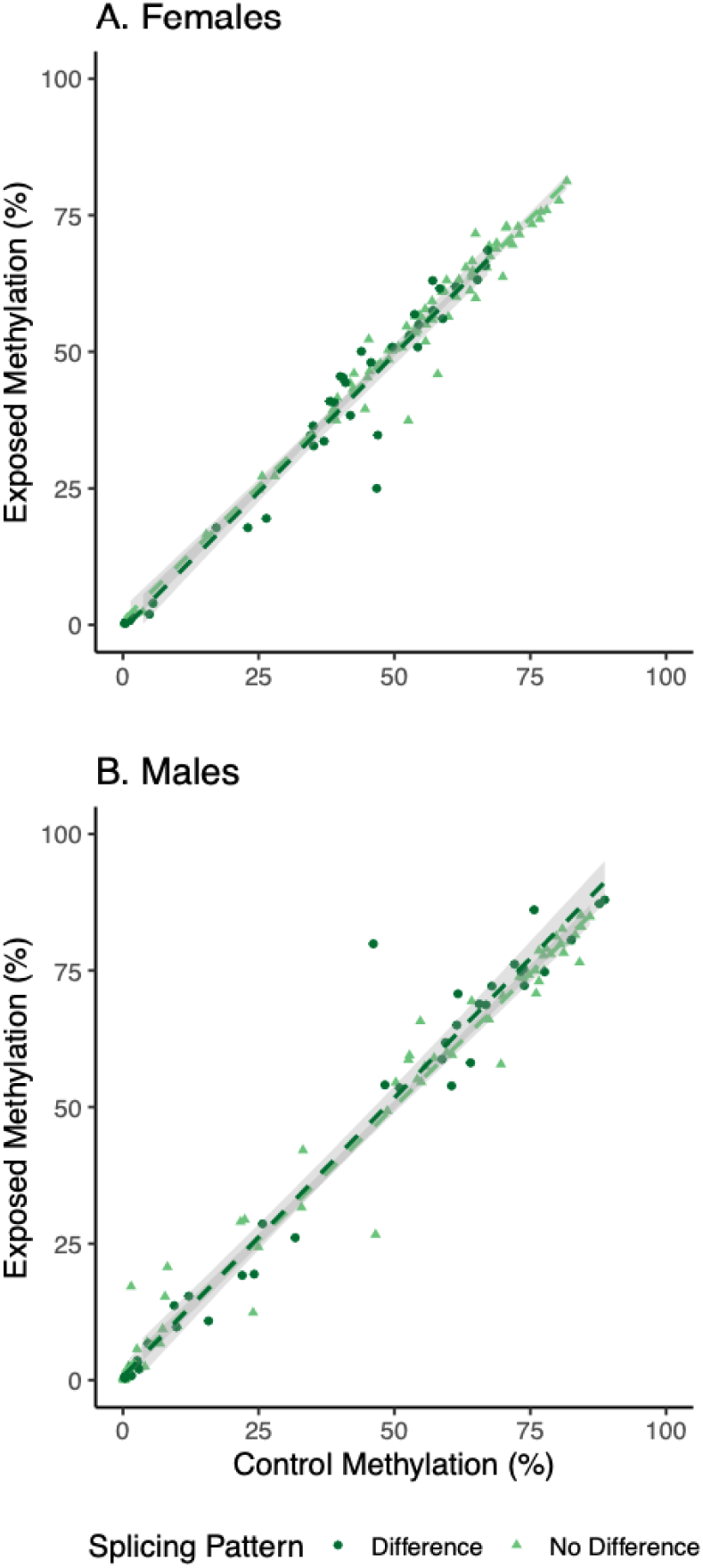
Average gene body methylation in control and exposed samples for genes that were alternatively spliced (difference) and not alternatively spliced (no difference) based on elevated pCO_2_ exposure in A) females and B) males. Each point represents a gene that was one of the top 20 drivers of expression patterns in a PC. Regression lines with 95% confidence intervals are included for each gene category.

**Table 5.** Binomial GLM results evaluating the influence of gene body methylation and gene length on alternative splicing status for the top 20 genes from each ASCA PC. No explanatory variables were significant predictors of alternative splicing status.

| Predictor | Females |  |  | Males |  |  |
| --- | --- | --- | --- | --- | --- | --- |
| | $\beta$ Estimate | z-value | P-value | $\beta$ Estimate | z-value | P-value |
| Intercept | 4.80 | 1.67 | 0.09 | -2.91 | -1.25 | 0.21 |
| Gene Body Methylation in Control Samples | -0.02 | -0.44 | 0.65 | -0.04 | -1.03 | 0.30 |
| Gene Body Methylation in Exposed Samples | -0.02 | -0.37 | 0.71 | 0.04 | 1.14 | 0.25 |
| $\log_{10}(\text{Gene Length})$ | -0.85 | -1.26 | 0.2 | 0.49 | 0.85 | 0.39 |

## Discussion

As elevated pCO_2_ continues to threaten marine invertebrates, it is critical to understand the role epigenetic and transcriptomic mechanisms play in stress resilience and the potential for environmental memory. Impacts on adult reproduction can influence intergenerational plasticity, so disentangling how various molecular mechanisms influence reproductive tissue is key for understanding offspring responses to stress. For example, herbicide exposure [34], sea surface temperature [35], ocean acidification [36], and ocean acidification and warming [37] have elicited methylation patterns that have subsequently been inherited by offspring. McNally et al. [27] used adult oysters to produce offspring and understand intergenerational plasticity in elevated pCO_2_ conditions. After three days of exposure, larvae from parents exposed to elevated pCO_2_ had higher shell growth rates and larger shell sizes, and these effects were pronounced when larvae were also reared in elevated pCO_2_ conditions [27]. Interestingly, there was no impact of elevated pCO_2_ on egg size or shape [27]. Since maternal provisioning may not have been responsible for larval phenotypes, exploring methylation and gene expression responses in gonadal tissues of those same adult oysters can elucidate regulatory mechanisms underlying observed carryover effects. By better understanding these mechanisms, we can better predict how environmental stress will influence intergenerational responses, and therefore ecosystem outcomes.

Given that the functional role of DNA methylation in marine invertebrates remains unclear [33], we leveraged whole genome, transcript-level data to conduct the first extensive assessment of DNA methylation-gene activity relationships in response to elevated pCO_2_ in *Crassostrea* spp. gonad tissues to our knowledge. We found that DNA methylation was significantly correlated with gene expression variability, or transcriptional noise, in a sex-specific manner in response to elevated pCO_2_. These findings align with the growing consensus that gene body methylation may not have a direct regulatory role in gene-level expression, but instead functions to broadly reduce transcriptional variation across the genome [33,36,40,55,56]. Additionally, differences in gene activity and methylation patterns between female and male oysters highlight the need to examine how OA may affect reproduction differently in females and males [21], and underscore the cell- and tissue-specificity of DNA methylation [57,58].

### Methylation does not impact gene-level responses in gonads to elevated pCO_2_

In this study there was no substantial impact of elevated pCO_2_ on gene-level expression in female gonads and male sperm. This is not surprising when considering a similar study on elevated pCO_2_ on gene expression in *C. virginica* mantle tissue [42]. Additionally, the heterogeneity of cell types (eggs, connective tissue) in female samples could have increased expression variability and reduced the ability to detect DEG. There are several OA studies that do see an impact on gene expression [20], with the predominant functions of influenced genes associated with stress response, acid-base regulation, metabolic processes, and calcification. While we did not see substantial gene-level impacts, we did see significant changes to transcript-level activity — changes in the maximum number of transcripts expressed per gene, shifts in the predominant transcript expressed, and patterns of alternative splicing — and to DNA methylation. These changes were associated with biological processes such as reproduction (*ex.* gamete development, regulation of female receptivity, sperm motility) various signaling pathways, and protein ubiquitination. These data suggest the importance of these molecular mechanisms in oyster response to elevated pCO_2_ conditions.

Beyond understanding the response of oyster reproductive tissue to elevated pCO_2_, a primary impetus for this work was to address hypotheses surrounding the role of methylation in environmental response and gene expression regulation. Following mammalian studies, it was previously considered that environmentally-sensitive changes in methylation drive differential expressed genes on a gene-by-gene basis in marine invertebrates [59,60]. However, to date, numerous molluscan studies have failed to find any significant relationships between differential DNA methylation and differential gene expression [42,45,49], or have found minimal overlaps in differentially expressed genes and differentially methylated regions [48,61]. As described above, we could not examine the association between differential methylation and differential gene expression since we did not observe any of the latter. We also did not see a concordant change in gene expression at the gene level where differential methylation occurred. Taken together, our data suggests that changes in methylation within a gene do not drive concurrent changes in average gene expression in gonad tissue, which is similar to what has been previously reported in molluscan somatic tissue [42,45,49].

### Methylation regulates gene activity differently in female and male reproductive tissue

A critical regulatory role for DNA methylation is the reduction of gene expression variability, or transcriptional noise. We speculate that this genome-wide role may be more important than direct modulation of gene expression. This study found an inverse relationship between methylation and transcriptional noise: genes with higher levels of methylation had lower expression variation (**Figure 5**). The relationship found in this study is consistent with previous research in molluscan somatic tissues [42,45,49,62] and across invertebrate taxa [33,55,56]. We also demonstrate that methylation variability is positively correlated with transcriptional noise (**Figure 5**). In other words, larger changes in gene body methylation are associated with higher expression variability. The general relationships described above were also consistent in response to elevated pCO_2_, further supporting the idea that methylation has a broad regulatory role in response to stress. In this study, we show that these responses are also seen in gonad tissues. We also identified a sex-specific pattern: elevated pCO_2_ decreases transcriptional noise in female gonads but increases noise in male sperm with respect to control conditions (**Figure 5**). The difference in methylation-noise relationships between sexes could suggest that sperm cells require additional methylation to achieve similar levels of transcriptional noise reductions to females. As egg quality is essential to successful reproduction, reducing transcriptional noise may be necessary to maintain egg quality, while gene expression variability may not be maladaptive for sperm. Additionally, eggs contain higher levels of RNA than sperm [63], so modulating transcriptional noise may be more important in eggs. Given the small effect size of pCO_2_ treatment on transcriptional noise in males, future work should investigate the phenotypic implications of these shifts. Combined with the subtle gene expression differences between control and OA-exposed oysters, our work suggests that reductions in general gene variability are an integral component of OA response in gonad tissues of *C. virginica*.

Additional differences in female and male reproductive tissue responses to elevated pCO_2_ underscore the importance of disentangling biological and environmental effects on gene expression and DNA methylation. It is well-known that OA can have sex-specific impacts on physiology [21], and previous work in *C. gigas* has documented distinct gene expression patterns associated with oogenesis and spermatogenesis [53]. We extend this work by demonstrating contrasting OA effects between female gonads and male sperm. We found that sperm had more genes with higher number of transcripts expressed in elevated pCO_2_, as opposed to female gonad tissue that had more genes with a smaller number of transcripts expressed in the same conditions (**Table 1**). Even though OA-responsive patterns of alternative splicing were similar between sexes, the top 20 gene drivers of those patterns differed and were involved in sex-specific reproductive processes. For example, protein UFMylation genes were alternatively spliced in female gonad tissue, while genes involved in regulation of metallopeptidase activity were alternatively spliced in male sperm. Protein UFMylation is involved in broader protein ubiquitination processes, which are conserved across oyster responses to pCO_2_ [11,43] and characteristic of female gonad methylome responses [46]. Metallopeptidases are important for sperm motility in humans [64,65], and have been found to increase in expression in *C. gigas* during spermatogenesis [66].

Differences in female and male reproductive tissue responses extend to methylation as well. Sperm had a more dramatic methylation response to elevated pCO_2_, with 2,916 DML compared to 89 in female gonad tissue oysters, even though a similar number of CpGs with 10x coverage were detected in both sexes. Sex-specific differences in methylation [50–52] are well-documented in *C. gigas*. Male *C. gigas* have higher levels of gene body methylation than females in both somatic [52] and reproductive tissue [50,51]. Our results confirm that these patterns are present in *C. virginica* gonads as well. These gene expression and methylation differences may be attributed to cell type: oocytes typically have higher levels of maternal RNA as opposed to sperm cells with more DNA [63]. Enrichment of processes such as oocyte development in females and spermatogenesis in males in genes with altered activity or with DML align with the specific biological roles of these cell types. In our study, female samples were likely a mix of oocytes and connective tissue, which further complicates the observed patterns. Previous studies have documented environmentally-sensitive methylation in response to elevated pCO_2_ in molluscan reproductive tissue [44,46], but none examine sex-specific patterns or have data for both sexes. As the role of methylation is cell-, tissue-, and sex-specific [57,58], future investigations should compare molecular and phenotypic responses to elevated pCO_2_ between tissues and sexes when appropriate.

### Methylation alters some aspects of transcript-level gene activity in gonad tissues

Although there is limited evidence for DNA methylation driving differential gene expression in molluscs, it is possible that methylation may change transcriptional opportunities. Sparse methylation may allow for exon skipping or access to alternative start sites, thereby modifying gene expression and phenotype [32,33,67]. Using differential exon expression as a proxy for alternative splicing, we observed that elevated pCO_2_ changes exon expression patterns (**Figures 1-2**). However, environmentally-responsive splicing accounted for a small proportion for the total splicing variance. There were minimal overlaps with DML and alternatively spliced genes, and gene body methylation did not significantly influence splicing (**Table 5**). This contrasts with work in *C. gigas* and upwelling-exposed *Strongylocentrotus purpuratus* urchin larvae showing that high levels of methylation in exons [68,69] and/or introns [69] are associated with alternative splicing. In *C. gigas*, alternative splicing was explored in one gene (CGI_10021620) with higher methylation and gene expression in mantle tissues as opposed to male gametes [68]. The authors conclude that lower levels of methylation in male gametes may have opened access to an alternative stop codon, producing a shorter protein product when compared to the product from mantle tissues [68]. Since *C. gigas* were not exposed to any stressors in this previous study, the role of methylation in regulating alternative splicing could be distinct for cell type differentiation versus environmental response. Significant methylation regulation of alternative splicing was found in *S. purpuratus* larvae from female urchins exposed to upwelling, but only when considering transcriptional start site accessibility [69]. It is possible that incorporating additional epigenetic mechanisms like chromatin state, or utilizing long-read sequencing methods for gene expression data, would elucidate connections between methylation and alternative splicing in *C. virginica*.

The lack of correlation between methylation and alternative splicing in our study could indicate that these molecular mechanisms are not intrinsically linked in *C. virginica*. In both sexes, we observed distinct biological functions associated with alternatively spliced genes and genes containing DML. For example, in female oysters alternatively spliced genes were involved in metabolic and developmental processes such as the glycolysis pathway and DNA ligation, while genes with DML had prominent roles in apoptosis, reproductive processes, and cell organization and biogenesis. The latter processes have been previously implicated in oyster response to elevated pCO_2_ [11,43], including studies examining gonad methylome responses [44,46]. Another potential explanation is that methylation is decoupled from transcript activity in gametes. The majority of RNA in oocytes are acquired during vitellogenesis, which may not reflect gene expression responses to OA [63]. On the other hand, sperm cells have lower RNA content and are not as transcriptionally active [63]. Therefore, it is possible that DNA methylation and alternative splicing have separate, but essential, roles in moderating reproductive tissue response to OA.

Although methylation and alternative splicing patterns were not correlated, the influence of gene body methylation on other aspects of gene activity suggests a potential regulatory role. In male oysters, methylation was significantly associated with changes in the maximum number of transcripts expressed per gene, and was marginally associated with shifts in the predominant transcripts (**Tables 2-3**). Interestingly, genes both containing DML and exhibiting alterations to gene activity had similar enriched biological processes involved in signal transduction and cell organization and biogenesis. These commonalities further support the idea methylation may have a role in gene activity beyond regulating alternative splicing patterns for specific pathways.

### Conclusion

Elucidating the relationship between epigenetic and transcriptomic mechanisms will allow for a holistic understanding of phenotypic responses to climate stressors. We highlight the importance of transcript-level data for parsing the relationship between these mechanisms. Future research should integrate transcript-level data along with exon- and intron-specific methylation and expression analyses to further clarify the relationship between methylation and gene activity within and across generations. As DNA methylation is a potential mechanism for intergenerational carryover effects [34–37], it is critical to assess environmentally-responsive methylation in adult reproductive tissue. Our study suggests that methylation has a genome-wide regulatory role, effectively maintaining gene expression homeostasis in reproductive tissues under elevated pCO_2_ by reducing transcriptional noise. Importantly, we found that the relationship between methylation and transcriptional noise was different between female reproductive tissue and sperm: male oysters required higher levels of methylation to achieve similar reductions in transcriptional noise when compared to female oysters. An accompanying study using the same adult *C. virginica* found no difference in female egg quality after OA exposure during reproductive conditioning, and no delays in early larval development [27]. Since this signal was observed even with significant genotype effects [27], it suggests that epigenetic maintenance of reproduction may also confer intergenerational resilience to elevated pCO_2_. We demonstrate that OA conditions lead to molecular changes in oyster gonads, and future work should evaluate specific connections between these changes and offspring fitness to understand whether inheritance of environmental memory can occur.

## Materials and Methods

### Experimental Design and Seawater Chemistry Analysis

The experimental design and seawater chemistry manipulations have been published by McNally et al. [27]. Briefly, adult *C. virginica* (mean ± SD shell length: 7.92 ± 1.80 cm) were acclimated to ambient pCO_2_ conditions (mean pCO_2_ ± SD = 632 ± 64 ppm) for one week prior to random placement in experimental pCO_2_ conditions (either control (572 ± 107 ppm) or elevated pCO_2_ (2827 ± 360 ppm)) during reproductive conditioning at 20°C. The elevated pCO_2_ treatment corresponds with seawater conditions undersaturated with aragonite, and is consistent with observations in estuarine ecosystems that oysters inhabit [42,70]. Additionally, the elevated pCO_2_ treatment was chosen to increase the inferential and statistical power of the study [71].

Four replicate 42-L tanks containing 10 oysters each were used for each pCO_2_ condition. Oysters were exposed to experimental conditions for 30 days, which has shown to elicit differences in *C. virginica* gonad methylation [44]. A flow-through ocean acidification array was used to control temperature, salinity, and pCO_2_ with temperature, salinity, and pH_T_ recorded three times per week during the experimental period. Seawater samples were collected during the first and third weeks of the experiment for dissolved inorganic carbon and total alkalinity measurements. Oysters were fed in accordance with best practices from Helm and Bourne [72]. No adult mortalities or significant random effects of adult tank on egg quality were reported [27]. At the end of the 30-day exposure, gonad samples from adult oysters were visually inspected under a microscope to determine sex and maturity, with sperm samples also checked for motility, targeting at least eight mature females and eight mature males per treatment [27]. Gonad tissues were collected from 16 female oysters (eight from three replicate control tanks and eight from two replicate elevated pCO_2_ tanks), and motile sperm from ten male oysters (four control and six exposed to elevated pCO_2_), with at least one individual from each replicate tank, for WGBS and RNA-Seq.

### Nucleic Acid Extraction and Library Preparation

Simultaneous DNA and RNA extraction was conducted for female gonad and sperm samples using the Zymo Quick-DNA/RNA Microprep Plus kit (Cat # D7005). A total of 10-15 mg of frozen gonad tissue and 50 µL of sperm in seawater solution were used as inputs for females and males, respectively. Manufacturer’s instructions were used, with the following exceptions. After addition of 300 µL DNA/RNA Shield (1X), 30 µL PK Digestion Buffer, and 15 µL Proteinase K, female samples were incubated at 55°C overnight. Four volumes of DNA/RNA Lysis Buffer (200 µL) were added to sperm samples. All centrifugation steps were conducted at 16,000 rcf. DNase I digestion was performed for RNA. Concentration of eluted DNA and RNA were verified with Qubit dsDNA High Sensitivity or RNA High Sensitivity assays (Thermo Fisher Scientific), respectively, and quality confirmed on a BioAnalyzer.

All samples were submitted to ZymoResearch for library preparation and sequencing. WGBS libraries were created from 100 ng of DNA with the Zymo-Seq WGBS Library Kit (Cat#: D5465) following manufacturer’s protocol and PCR performed with Illumina Unique Dual Indices. Library quality was checked with an Agilent 2200 TapeStation. Libraries were sequenced on an Illumina NovaSeq instrument to generate 150 bp paired-end reads. Prior to sequencing, total RNA-Seq libraries were created using 250 ng of RNA. For all samples, rRNA was removed following methods described in Bogdanova et al. [73] with some modifications.

Libraries were constructed using the Zymo-Seq RiboFree Total RNA Library Prep Kit (Cat # R3000) according to the manufacturer’s instructions. Libraries were sequenced on an Illumina NovaSeq to generate at least 30 million read pairs for 150 bp paired-end sequences per sample. Resulting sequences were submitted to the NCBI Sequencing Read Archive (SRA) under BioProject accession number PRJNA1089235.

### Genome Information and Feature Tracks

The *C. virginica* genome was used for several analyses [74]. Gene, coding sequence, exon, and lncRNA tracks were provided by the *C. virginica* genome NCBI RefSeq annotation (RefSeq: GCF_002022765.2) on NCBI [74]. These tracks were used to generate remaining feature tracks with BEDtools v2.26.0 [75]. The complement of the exon track, or non-coding sequences, was created with BEDtools `complementBed` v.2.26.0. Overlaps between these non-coding sequences and genes — introns — were obtained using `intersectBed`.

Untranslated regions of exons were identified by subtracting the coding sequence track from the exon track with `subtractBed`. Upstream flanking regions, or putative promoters, were defined as the 1000 bp upstream of a transcription start site. These flanks were created by adding 1000 bp upstream of genes with `flankBed`, taking strandedness into account. Other genes that overlapped with these flanks were removed using `subtractBed`. Flanks 1000 bp downstream of genes were created as well. To identify intergenic regions, the complement of the gene track was created with complementBed, then `subtractBed` was used to remove flanking regions from the intergenic region track. Transposable element information from RepeatMasker was obtained from the NCBI RefSeq annotation [76,77]. CG motifs in the *C. virginica* genome were identified using the `fuzznuc` function from EMBOSS. The number of CG motifs in a given track were obtained using `intersectBed` between the CG motifs and a feature track. The genome features are available along with corresponding code [78].

### Genetic Variation

SNPs were identified in the BS-seq data using the Nextflow [79] EpiDiverse/snp pipeline [80] to characterize genetic differences between samples. Sorted, deduplicated BAM files, generated through methylation analysis with Bismark [81], were used as inputs, along with the National Center for Biotechnology Information (NCBI) *C. virginica* genome FastA (`GCF_002022765.2_C_virginica-3.0_genomic.fa`) provided as the reference. The options `--variants` and `--clusters` were provided when running the EpiDiverse/snp pipeline with default settings (`--min-coverage 0`, `--min-repeat-entropy 1`). After variant identification, NgsRelate was used to estimate pairwise relatedness using VCF output from EpiDiverse/snp [82]. Variants were filtered to include only bi-allelic sites (`--min-alleles 2--max-alleles 2`) and minor allele counts of at least two (`--mac 2`). Variants were included if data was missing for more than 50% of the samples (`--max.missing 0.5`).

Recent studies have shown that genetic population structure can influence methylation [83]. In order to determine if genetic variation had a significant impact on gene expression and methylation, we first performed correlation analyses to determine whether genetic distance correlated with methylation and gene expression distance between individuals. In the case of a significant correlation, genetic distance would need to be included in downstream model analyses of differential methylation and gene expression. To perform these correlations, we generated euclidean distance matrices of genetic data, expression data, and DNA methylation data. For genetic data, SNPs were derived from the Nextflow EpiDiverse SNP Pipeline as described above. Gene expression matrices were derived from `DESeq2` output [84]. DNA methylation data were obtained from `methylkit` output using a 10x coverage minimum threshold. For analysis, Mantel tests (`mantel.rtest`) in R were used with 99 replicates. We visualized these relationships using Pearson correlation matrices. Corresponding code is available for all genetic variation analyses [78].

### Gene Activity

#### Gene and Transcript Expression

FastQ files were quality trimmed using `fastp` [85] default settings for paired end adapter trimming, followed by the removal of the last 20 bases from the five prime end of all reads. These bases were removed because the FastQC “Per Base Sequence Content” assessment showed these exhibited less uniform distribution of nucleotides than the remainder of the read length. Trimmed reads were then visualized with `MultiQC` [86]. To identify isoforms, the *C. virginica* genome (NCBI accession: GCF 002022765.2) was indexed using HISAT2 [87]. A pipeline detailed by Pertea et al. [88] was used as a guide to identify isoforms and analyze differential expression. Trimmed reads were aligned to the genome using HISAT2 (v2.1.0), followed by StringTie [89] to identify isoforms and generate files for downstream analysis using the R [90] packages `ballgown` [91] and `tidyverse` [92]. Differentially expressed genes (DEG) and differentially expressed transcripts (DET) (p-value and q-value < 0.05) were identified between pCO_2_ treatments within a given sex using `ballgown` [91], with ‘`FPKM`’ as the expression measurement.

Elevated pCO_2_ may influence finer-scale expression processes. To explore this, we quantified differences in the maximum number of transcripts expressed in each gene for each sex separately. The maximum number of transcripts for each gene was identified across all samples. The difference in maximum number of transcripts expressed in each gene was calculated with respect to the elevated pCO_2_-exposed samples. Positive differences signify more unique transcripts expressed in elevated pCO_2_ conditions, while the opposite holds true for negative differences.

An enrichment test with `topGO` [93] was conducted to understand if certain biological processes were overrepresented in genes with differences in the maximum number of transcripts expressed. First, a gene ID-to-GO term database (`geneID2GO`) was created for manual GO term annotation. Each line of the database contained gene ID in one column, and all corresponding GO terms in another. The list of gene IDs from this database was used as the gene universe for enrichment. Two separate lists were compiled for genes of interest: one for genes with increased transcript counts, and another for genes with decreased transcript counts due to elevated pCO_2_ exposure. A `topGO` object was generated for each sex- and direction-specific list of genes and biological process GO terms, with GO term annotation performed using the `geneID2GO` database. A Fisher’s exact test was used to identify GO terms significantly enriched with respect to the gene background (*P*-value < 0.01). Corresponding code is available for all gene activity analyses [78].

### Predominant Transcript Identification

In addition to quantifying transcript expression, we examined shifts in the predominant transcript — the transcript with the highest mean level of expression in an experimental group — due to low pH exposure within male and female samples separately. Gene expression data from `ballgown` was filtered to retain genes with multiple transcripts where gene expression was greater than zero. Differences in the identity of the predominant transcript were determined for genes remaining after filtering. An enrichment test with `topGO` [93] was used to identify significantly enriched biological process GO terms (*P*-value < 0.01) in genes where the predominant transcript changed using similar methods as described above (see *Gene and Transcript Expression*).

### Alternative Splicing

The final way we characterized gene activity was through an alternative splicing analysis. We hypothesized that OA exposure would induce different patterns of exon expression, or alternative splicing. To characterize these patterns, expression of exons 2-6 were calculated as fold change values relative to expression of exon 1 for every gene in each sample. To do this, `bedtools::coverage` was used with BAM alignment files from HISAT2 and the genome feature file. A minimum read count coverage of 10x was used. These values were generated for females and males separately. We then used ANOVA simultaneous component analysis (ASCA) using the `ALASCA` package (v1.0.14) in R [94] to perform longitudinal multivariate analysis across exon position. ASCA is useful for the analysis of both fixed and random effects on high-dimensional multivariate data when other multivariate ANOVA analyses (i.e., MANOVA) are not able to include more variables than there are observations [54]. Each sex-specific ASCA model analyzed multivariate expression as a function of exon position, treatment, and their interaction, with the sample as a random effect to account for repeated measures:

~~~
expression relative to exon 1 ∼ exon position * treatment + (1|sample)
~~~

Genes that were detected in all samples were included (11,270 genes in females and 12,645 genes in males). The model was validated with bootstrapping with replacement for 100 iterations [95]. We extracted and plotted principal components (PC) that explained greater 1% of the variance to visualize expression patterns across exons. We categorized PC as those that either 1) show no difference in exon expression patterns between treatments or 2) show differences between treatments. For each PC, we extracted the top 20 genes with the highest PC scores. These gene lists were combined into the two previously described categories for downstream analyses. For these top genes, the number of genes with changes in the maximum number of transcripts expressed were quantified for females and males.

A functional enrichment analysis was conducted with `topGO` [93] to understand if alternatively spliced genes contained any overrepresented biological processes. Enrichment was conducted using similar methods as described above (see *Gene and Transcript Expression*). All genes included in the ASCA analysis were used as the gene background for this analysis. Biological process GO terms with *P*-values < 0.01 were considered significant.

### DNA Methylation Analysis

#### CpG locus-level Methylation Quantification

Quality-trimmed sequence reads were aligned to the *C. virginica* genome using Bismark v.0.22.3 using `-score_min L,0,-0.6` to dictate alignment specificity. In order to characterize CpG methylation for individual samples, bedGraph files were generated based on Bismark `coverage2cytosine` output, where CpG data from each strand was merged (`--merge_CpG`), filtered to retain CpG loci with at least 10x coverage, and sorted. Since C->T SNPs can be interpreted as unmethylated cytosines, they were identified in bedGraphs using default BS-Snper settings [96], then the methylation values of those C->T SNPs were changed to 0.

Sex-specific baseline methylation landscapes were characterized using methylation information for CpG loci with at least 10x coverage in a sample. Individual bedGraph files (described above) were combined using `bedtools::unionBedGraphs` v2.30.0 [75] to create sex-specific union bedGraphs. Percent methylation was averaged across all samples for each CpG locus. Highly (> 50%), moderately (10-50%), and lowly (≤ 10%) methylated CpGs were identified in sex-specific union bedGraphs. Overlaps between highly methylated CpGs and various genome features were characterized using `bedtools::intersectBed`. A chi-squared contingency test (`chi.sq`) was used to determine if the distribution of highly methylated CpGs was significantly different from the distribution of all CpGs in the *C. virginica* genome, with *P*-values < 0.05 indicating a significant difference.

Deduplicated BAM alignment files from Bismark were used for sex-specific differential methylation analysis with `methylKit` [97]. The threshold for comparison was a minimum of 10x coverage (`lo.count = 10`) and maximum coverage set at 98% (`hi.perc=98`), using the destranded setting (`destrand = TRUE`). Putative differentially methylated loci were determined based on a 50% difference in comparison group and q-value threshold of 0.01.

Finally, C->T SNPs were removed from putative female and male DML lists using `bedtools::subtractBed`. Genomic location of female and male DML were determined by overlapping with annotated features using `bedtools::intersectBed`. A chi-squared contingency test was used to test the null hypothesis of no association between methylation and specific genome features using all CpGs with 10x data and DML for females and males separately. A *P*-value < 0.05 signified a significantly different proportion of DML in a specific genome feature when compared to 10x CpG data. An enrichment test with `topGO` [93] was used to identify significantly enriched biological process GO terms (*P*-value < 0.01) in genes containing DML The gene universe for the enrichment was a sex-specific list of genes containing 10x CpGs. The enrichment was conducted using similar methods as described above (see *Gene and Transcript Expression*).

### Gene-level Methylation Quantification

Gene level methylation was determined by taking the mean value of methylation at loci within a gene where coverage was at least 10x. Specifically, bedgraph files generated as part of the Bismark alignment described above were used in conjunction with `bedtools::intersectBed` to extract CpG loci for each sample, for each gene.

### Methylation Influence on Gene Activity

To broadly examine methylation and gene expression signatures of OA treatment for each sex, we used a similar approach to the genetic variation analysis. We used Monte Carlo tests (`mantel.rtest`) to calculate correlations between bray-curtis distance matrices derived from gene methylation and expression datasets. Separate subsequent analyses were conducted to examine if methylation influenced specific aspects of gene activity.

### Influence of Methylation on Gene and Transcript Expression

The first set of analyses were used to determine the relationship between gene methylation and expression. First, the number of DML in DEG and DET were quantified. Next, we investigated if gene methylation influenced differences in the maximum number of transcripts expressed between control and exposed samples using a binomial GLM (`glm`). For each sex separately, we examined if a change in the maximum number of transcripts expressed due to OA exposure (“success”) could be explained by change in gene body methylation, number of DML in a gene, change in gene expression, or gene length:

~~~
occurrence of a change in the maximum number of expressed transcripts
∼ change gene methylation + number of DML in the gene + log2(change in gene expression + 1) + log10(gene length)
~~~

Change in gene body methylation was calculated by subtracting average gene body methylation in the control treatment from the OA treatment. A similar calculation was done to obtain change in gene expression. Explanatory variables were considered significant if *P*-values < 0.05. Corresponding code is available for all gene activity analyses [78].

### Influence of Methylation on Predominant Transcript Shifts

The second set of analyses examined if previously-identified transcript shifts were due in part to gene methylation. Similar sex-specific binomial GLM (`glm`) to those described above (see *Influence of Methylation on Gene and Transcript Expression*) were constructed to examine if occurrence of a predominant transcript shift due to OA exposure could be explained by change in gene body methylation, number of DML in a gene, change in gene expression, or gene length:

~~~
occurrence of predominant transcript shift ∼ change gene methylation
+ number of DML in the gene + log2(change in gene expression + 1) + log10(gene length)
~~~

Predictors were significant if *P*-values < 0.05.

### Transcriptional Noise Analysis

Following Wu et al. [98], transcriptional noise, or the CV of gene expression, was used as a response variable in a linear model (`lm`) with average gene methylation, average gene expression, CV of gene methylation, gene length, OA treatment, and the interaction between treatment and methylation or expression:

~~~
log_10_(CV of gene expression) ∼ gene methylation + log_2_(gene expression
+ 1)^2^ + log_2_(gene expression) + log_10_(CV of gene methylation) + log_10_(gene length) + treatment + treatment*gene methylation + treatment*log_2_(gene expression + 1)^2^ + treatment*log_2_(gene expression + 1)
~~~

The polynomial terms for gene expression were included after initial data visualization, and represent a relationship between gene expression and transcriptional noise where gene expression variability is highest at very low or high levels of gene expression. A backwards deletion approach was used for model construction, with explanatory variables removed from the model based on a *P*-value < 0.05 threshold. Competing models were evaluated with the Akaike information criterion (AIC). The best model fit was defined by the lowest AIC and all input variables having a *P*-value < 0.05. Residuals of the best fit model were evaluated for normality and homoscedasticity. Coefficients of determination, or partial R^2^, were calculated for all significant explanatory variables. To calculate partial R^2^, the sum of squares from `anova` output was divided by the total sum of squares for all input variables and residuals.

### Influence of Methylation on Alternative Splicing

Finally, methylation’s influence on gene activity was evaluated by understanding methylation patterns in alternatively spliced genes. The number of DML were counted in genes that were and were not alternatively spliced. A sex-specific binomial GLM (`glm`) was used to determine if average gene body methylation in control or exposed samples and gene length were significant predictors of alternative splicing:

~~~
whether or not the gene was alternatively spliced ∼ average gene body methylation in control samples + average gene body methylation in exposed samples + log10(gene length)
Terms were considered significant predictors of alternative splicing if *P*-values < 0.05.
~~~

## Acknowledgements

This work was facilitated through the use of advanced computational, storage, and networking infrastructure provided by the Hyak supercomputer system at the University of Washington. This work was funded by the National Science Foundation awards 1634167 to SBR and 1635423 to KEL. We are grateful to Grace Crandall for assistance with DNA and RNA extractions and to Drs. Shayle Matsuda, Katherine Silliman, Laura H. Spencer, and Samuel N. Bogan for insight on manuscript ideation and earlier drafts of this manuscript.

## Data Availability

Raw WGBS and RNA-Seq data can be accessed at the NCBI Sequence Read Archive under BioProject accession number PRJNA1089235. Additional data, genome feature tracks, scripts, and supplementary materials are available in the *C.virginica* methylation and gene expression repository on Open Science Framework (https://osf.io/xuy2f/) and on Github (https://github.com/sr320/ceabigr) [78].

## Author Contributions

AD-W and KEL conceived and ran the experiment. YRV performed DNA and RNA extractions. All authors contributed to initial analyses. YRV, ASH, SJW, and SBR conducted final analyses. AD-W and KEL contributed to analysis. YRV, ASH, SJW, and SBR wrote the initial manuscript draft. All authors reviewed and approved the manuscript.

