## Supplementary Files for "DNA methylation correlates with transcriptional noise in response to elevated pCO_2_ in the eastern oyster (*Crassostrea virginica*)"

All supplementary files are at https://github.com/sr320/ceabigr/tree/main/supplemental-files

**Supplementary File S1**. Spearman correlation matrix of **A)** gene expression distance and genetic distance between individuals; **middle)** methylation distance and genetic distance between individuals; and **right)** methylation distance and gene expression distance between individuals. Distance matrices for each response were generated using euclidean distance in the dist() function. Blue scale dots indicate significant positive correlations and red scale dots indicate significant negative correlations with color indicating strength of correlation. Numbers in rows and columns indicate individuals (M indicates males, F indicates females). Positive correlations between genetic distance and methylation of gene expression indicate that individuals more closely related to each other have greater similarity in methylation gene expression. Positive correlations between gene expression and methylation indicate that individuals with greater similarity in methylation are also more similar in their gene expression. Relationships were tested with Mantel Tests.
https://github.com/sr320/ceabigr/blob/main/supplemental-files/S1-genetic_correlations_figure.pdf

**Supplementary File S2**. Volcano plots showing transcript and gene expression in females. A single transcript was identified in female controls as being differentially expressed (p-value and q-value <= 0.05).
https://github.com/sr320/ceabigr/blob/main/supplemental-files/S2-faceted-volcano_plots-fmcoe-DETs-DEGs.png

**Supplementary File S3**. Gene expression and activity information for female and male samples. Information includes FPKM, number of unique transcripts expressed per gene in control and elevated $pCO_2$ conditions, and the identity of the predominant transcript in control and elevated $pCO_2$ conditions.
https://github.com/sr320/ceabigr/blob/main/supplemental-files/S3-01.01-fmcoe-max-predom-isos-gene_fpkm.csv

**Supplementary File S4**. Results from all enrichment tests conducted.
https://github.com/sr320/ceabigr/blob/main/supplemental-files/S15-all-enrichment-results.xlsx

**Supplementary File S5.** Variance explained by principal components identified by ANOVA-simultaneous components analysis (ASCA) in the analysis of gene expression variation across exon position (i.e., alternative splicing). Variance explained for each principal component shown for females (A) and males (B).
https://github.com/sr320/ceabigr/blob/main/supplemental-files/S4-ASCA-var-explained.pdf

**Supplementary File S6.** Principal components identified by ANOVA-simultaneous components analysis (ASCA) that describe patterns of alternative splicing that show no effect of treatment in females. (A) Pattern of exon expression described by PC1 with the top 10 genes associated with this pattern displayed in (B) ordered by PC loading score. (C) Pattern of exon expression described by PC2 with the top 10 genes associated with this pattern displayed in (D) ordered by PC loading score. (E) Pattern of exon expression described by PC3 with the top 10 genes associated with this pattern displayed in (F) ordered by PC loading score. (E) Pattern of exon expression described by PC4 with the top 10 genes associated with this pattern displayed in (F) ordered by PC loading score. In all plots, dark green indicates exposed treatment and light green indicates control treatment. Shading indicates 95% confidence intervals. Error bars indicate 95% confidence intervals with variables indicating genes.
https://github.com/sr320/ceabigr/blob/main/supplemental-files/S5-ASCA-female-PC1-4.pdf

**Supplementary File S7.** Principal components identified by ANOVA-simultaneous components analysis (ASCA) that describe patterns of alternative splicing that show no effect of treatment in males. (A) Pattern of exon expression described by PC1 with the top 10 genes associated with this pattern displayed in (B) ordered by PC loading score. (C) Pattern of exon expression described by PC2 with the top 10 genes associated with this pattern displayed in (D) ordered by PC loading score. (E) Pattern of exon expression described by PC3 with the top 10 genes associated with this pattern displayed in (F) ordered by PC loading score. (E) Pattern of exon expression described by PC4 with the top 10 genes associated with this pattern displayed in (F) ordered by PC loading score. In all plots, dark green indicates exposed treatment and light green indicates control treatment. Shading indicates 95% confidence intervals. Error bars indicate 95% confidence intervals with variables indicating genes.
https://github.com/sr320/ceabigr/blob/main/supplemental-files/S6-ASCA-male-PC1-4.pdf

**Supplementary File S8**. Genomic locations of all CpGs with 10x coverage, highly methylated CpGs, moderately methylated CpGs, and lowly methylated CpGs in A) females and B) males.
https://github.com/sr320/ceabigr/blob/main/supplemental-files/S7-meth-landscape-fig.pdf

**Supplementary File S9**. Chi-squared contingency test results comparing the location of highly methylated CpGs with all 10x CpGs in the *C. virginica* genome for female samples.
https://github.com/sr320/ceabigr/blob/main/supplemental-files/S8-fem-CpG-location-statResults.txt

**Supplementary File S10**. Chi-squared contingency test results comparing the location of highly methylated CpGs with all 10x CpGs in the *C. virginica* genome for male samples.
https://github.com/sr320/ceabigr/blob/main/supplemental-files/S9-male-CpG-location-statResults.txt

**Supplementary File S11**. Chi-squared contingency test results comparing the location of DML with highly methylated CpGs for female samples.
https://github.com/sr320/ceabigr/blob/main/supplemental-files/S10-CpG-location-statResults-Fem.txt

**Supplementary File S12**. Chi-squared contingency test results comparing the location of DML with highly methylated CpGs for male samples.
https://github.com/sr320/ceabigr/blob/main/supplemental-files/S11-CpG-location-statResults-Male.txt

**Supplementary File S13**. Number of DML in genes with differences in the maximum number of transcripts expressed or shifts in the predominant transcript for female and male samples.
https://github.com/sr320/ceabigr/blob/main/supplemental-files/S12-DML-counts-max-transcript-predominant-isoform.xlsx

**Supplementary File S14**. Change in average gene methylation (%) for genes that had an decrease in maximum transcript count (light green), increase in maximum transcript count (dark green), or no change (grey) in maximum transcript count due to OA exposure in **A)** female and **B)** male oysters. Change in methylation was a marginally significant predictor of whether or not there was a predominant transcript shift in males, not females.
https://github.com/sr320/ceabigr/blob/main/supplemental-files/S13-max-trans-change-meth-distribution.pdf

**Supplementary File S15**. Change in average gene methylation (%) for genes in **A)** female and **B)** male oysters with (dark green) or without (light green) a shift in the predominant transcript due to OA exposure. Change in methylation was a significant predictor of whether or not there was a predominant transcript shift in males, not females.
https://github.com/sr320/ceabigr/blob/main/supplemental-files/S14-pred-transcript-change-meth-distribution.pdf
